# Elevational changes in bacterial microbiota structure and diversity in an arthropod-disease vector

**DOI:** 10.1101/2021.06.02.446707

**Authors:** Tuomas Aivelo, Mélissa Lemoine, Barbara Tschirren

## Abstract

Environmental conditions change rapidly along elevational gradients and have been found to affect community composition in macroscopic taxa, with lower diversity typically observed at higher elevations. In contrast, microbial community responses to elevation are still poorly understood. Specifically, the effects of elevation on vector-associated microbiota have not been studied to date, even though the within-vector microbial community is known to influence vector competence for a range of zoonotic pathogens. Here we characterize the structure and diversity of the bacterial microbiota in an important zoonotic disease vector, the sheep tick *Ixodes ricinus*, along replicated elevational gradient (630 - 1673 masl) in the Swiss Alps. 16S rRNA sequencing of the whole within-tick bacterial microbiota of questing nymphs and adults revealed a decrease in Faith’s phylogenetic microbial alpha diversity with increasing elevation, while beta diversity analyses revealed a lower variation in microbial community composition at higher elevations. We also found a higher microbial diversity later in the season and significant differences in microbial diversity among tick life stages and sexes, with lowest microbial alpha diversity observed in adult females. No associations between tick genetic diversity and bacterial diversity were observed. Our study demonstrates systematic changes in tick bacterial microbiota diversity along elevational gradients. The observed patterns mirror diversity changes along elevational gradients typically observed in macroscopic taxa, and they highlight the key role of environmental factors in shaping within-host microbial communities in ectotherms.

## Introduction

Because of the rapid changes in abiotic conditions along elevational gradients, radical elevational species community turnovers are often observed within short geographical distances [1]. Indeed, temperature and oxygen pressure decrease, whereas ultraviolet radiation increases with increasing elevation [2]. Other abiotic factors, such as precipitation, wind velocity, seasonality, soil formation processes and disturbance typically also show systematic changes along elevation clines, but also vary by geographical context [2]. All of these abiotic factors affect species community composition and diversity along elevational gradients [1].

In macroscopic taxa, such as plants, invertebrates and mammals, biodiversity is typically found to either linearly decrease with increasing elevation or to show a hump-shaped pattern where diversity is highest at intermediate elevations [3]. In addition, community composition and the strength and direction of biotic interactions have been found to vary along elevational clines [4, 5]. Much less is known about how microbial communities change along elevational gradients, and the existing empirical studies suggest inconsistent elevation patterns that differ from patters observed in macroscopic taxa [e.g., 7–9].

If diversity gradients along elevational clines are different in microbes compared to macroscopic taxa, the underlying factors affecting these gradients are likely to differ as well. For macroscopic taxa, climatic variables seem to be the most important factors affecting diversity patterns [3], whereas for microbes, we do not yet have a good framework to understand trends in community composition [9]. To date, the best studied microbial communities are soil bacteria: their diversity seems to be mainly determined by the quality and composition of the soil, such as soil pH or carbon, without any systematic changes along elevational clines [6].

Compared to ‘free-living’ microbial communities, host-associated microbiota are shaped by an additional key factor: the identity, ecology, life history and quality of the host [10], which may affect microbial community composition along elevational gradients. However, patterns of diversity and community composition in host-associated microbiota does not seem to follow a consistent pattern either. For example, in pika (*Ochotona curzionae*) individuals living at higher elevation were found to have a higher alpha (i.e., the mean species diversity) and beta (i.e., heterogeneity in species composition) diversity in their gut microbial community compared to individuals living at lower elevation [11], whereas human skin microbiota shows a decrease in alpha diversity but an increase in beta diversity with increasing elevation [12].

Microbial communities of ectothermic hosts are expected to be most strongly affected by elevational gradients because ectotherms do not buffer ambient temperature as strongly as endothermic taxa. Furthermore, the microbiota of invertebrates is typically not stable [13]. Yet, changes in microbiota composition in invertebrate hosts along environmental gradients remain largely unexplored.

*Ixodes* ticks are common vectors for human pathogens including *Borrelia* sp. and *Rickettsia* sp., and recent studies suggest that the tick’s commensal microbiota composition affects the probability of harboring human pathogens [14]. Due to climate warming, *Ixodes ricinus* has expanded its distribution into both higher latitudes and higher elevations in many parts of Europe [15]. At the same time, the incidence of the diseases caused by tick-borne pathogens continues to increase in many regions [16]. The abundance and distribution of ticks are strongly influenced by temperature and other climatic variables: low winter temperatures increase tick mortality, whereas warmer temperatures during summer months lead to a faster life cycle and a longer activity period [17, 18].

While there is growing body of research on how changing environmental conditions might affect specific tick-associated microbes, specifically pathogens [19–21], no study has investigated how the structure of the commensal tick microbiota (or the microbiota of any other disease vector), changes along elevational clines and how this may indirectly affect pathogen prevalence and disease dynamics [21]. Abiotic variables may affect tick microbiota composition and diversity either directly through effects on microbial growth, competition and/or transmission [22], or indirectly through changed tick behavior or life history [17]. Furthermore, ticks quest in the undergrowth, attach to their vertebrate host and suck blood for a number of days. The tick microbiota is thus likely acquired from soil and plants but also from their host skin and blood [23], which all likely vary along elevational clines and may thus shape tick microbial communities.

Here we exploit the rapidly changing environmental conditions along elevational gradients in the Swiss Alps to quantify changes in tick microbiota diversity and community structure along elevational cline. Specifically, we test if tick bacterial microbiota diversity and community structure varies along elevational gradient, and if these patterns differ with season or across tick life stages / sexes. Furthermore, we analyzed how ecological processes influence community turnover by comparing phylogenetic relatedness within and between tick microbiota.

Based on previous findings, we predict systematic differences in tick bacterial microbiota composition along elevational gradients. Furthermore, because of differences in their behavior and ecology [17, 18], we predict differences in microbiota structure and diversity between tick life stages and sexes.

## Materials and methods

### Tick sampling and environmental data

We collected questing *Ixodes ricinus* ticks at three different locations in the Swiss Alps (Kanton Graubünden). Three sites per location were identified, one at low (630 - 732 m above sea level), one at medium (1 094 – 1 138 masl) and one at high (1 454 – 1 673 masl) elevation (Fig. 1; Table 1). At each site, tick sampling was performed thrice, once in June, once in July, and once in August 2014 by dragging a white blanket (1 m x 1 m) over the ground vegetation as described previously [24]. Coordinates and elevation were recorded for each site. We collected ticks from the blanket and stored them in 95% ethanol. Tick species and life stage were verified using a stereomicroscope.

**Figure 1:**
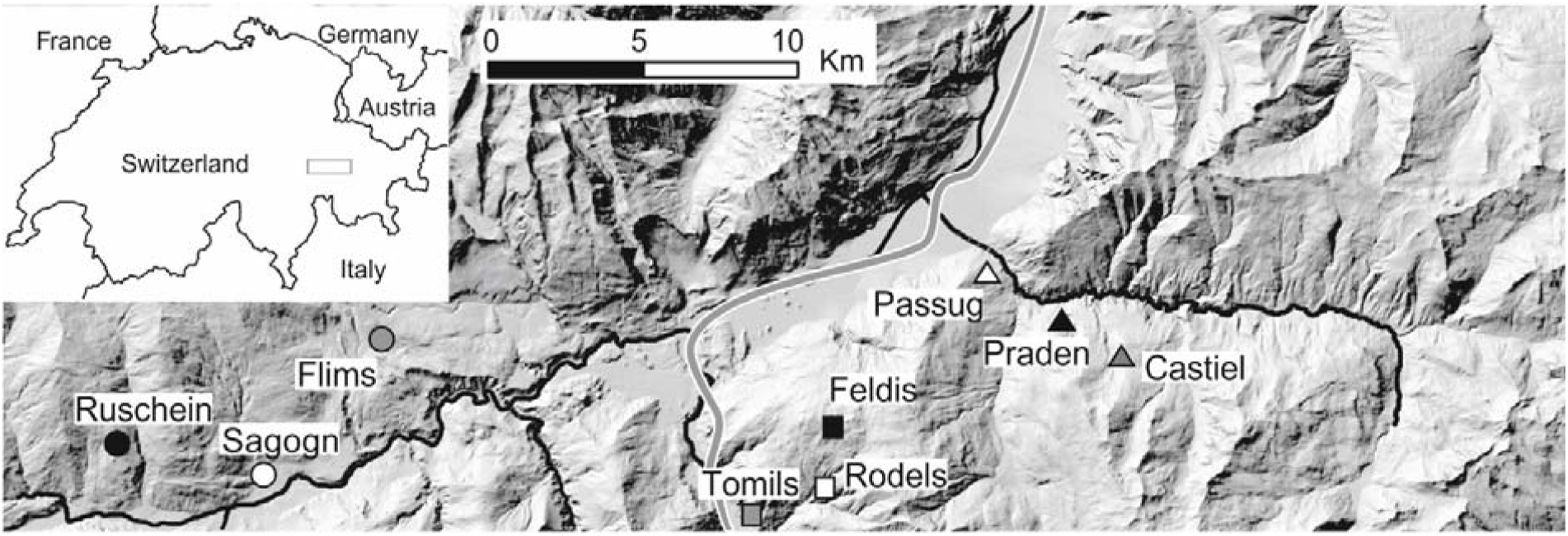
Location of tick sampling sites in the Swiss Alps. Different shapes (i.e., triangle, circle and square) represent the different locations, whereas colours represent the different elevations (white = low, grey = medium, black = high). Rivers are shown in black and motorway in grey. Figure is from Aivelo et al. (2020). Map data ©2019 Google, GeoBasis-DE/BKG.

**Table 1:**
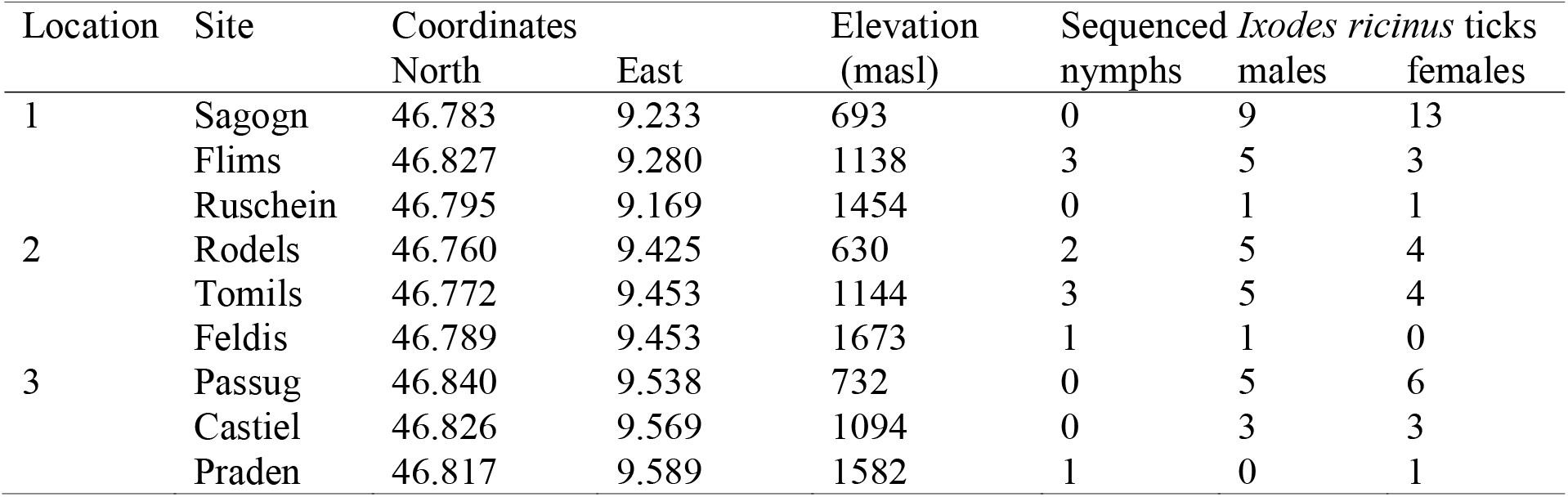
Tick sampling sites in the Swiss Alps.

### Tick microbiota

16S rRNA sequencing and tick bacterial OTU identification was performed as described previously [21]. In short, we randomly selected *I. ricinus* ticks from each sampling site (Table 1). DNA isolation and amplifications were performed in a laminar flow cabinet to avoid contaminations. Each tick was washed thrice with sterile water to remove surface bacteria before sterilizing it with 3% hydrogen peroxide. We cut ticks in half with a sterilized blade to facilitate DNA isolation and then used the DNeasy Blood & Tissue kit (Qiagen; Hilden, Germany) to extract DNA. We processed negative controls (N=5) alongside the tick samples.

We characterized tick bacterial community composition by sequencing the hypervariable V3-V4 region of the 16S gene by using the primers 515FB and 806RB [25] and prepared sequencing libraries following the Earth Microbiome 16S Illumina Amplicon protocol (see Supplementary materials S1 for details). To remove batch effects, we randomized samples and negative controls across two plates. The libraries were sequenced on Illumina MiSeq at the Functional Genomic Center Zurich with a target length of 250 bp following the manufacturer’s protocol. Sequences were analysed using the *mothur* pipeline with MiSeq standard operation procedures [26] with 99% similarity threshold for OTU clustering. Raw sequenced were deposited in the Sequence Read Archive under BioProject PRJNA506875. The metadata of the samples and their matching sequence accession numbers are deposited in FigShare (doi: <u>10.6084/m9.figshare.14540892.v1</u>), while code of all statistical analysis can be found in: https://github.com/aivelo/tick-biodiversity.

Using *mothur*, we purged unsuccessful contigs and preserved only contigs between 250 and 310 bp. The alignment was made against aligned SILVA bacterial references (release 128; https://www.arb-silva.de/documentation/release-128/). We used 99% similarity to determine OTUs and classified them using SILVA taxonomy. Only samples with > 500 reads and OTUs which were present in at least two samples were included in the analyses. We rarified the samples to 500 reads to account for variation in amplicon numbers.

### Statistical analysis

#### Bacterial alpha diversity

Bacterial taxonomic alpha diversity (inverse Simpson index; [27]) and Faith’s phylogenetic alpha diversity [28] were calculated with the R package *vegan* [29]. Additionally, we calculated two more phylogenetic alpha diversity indices: Nearest Relatedness Index, NRI, (equivalent to -1 standardized effect size of mean pairwise distances in communities, which estimates the average phylogenetic relatedness between all possible pairs of bacterial taxa within a tick) and Nearest Taxon Index, NTI, (−1 times standardized effect size of mean nearest taxon distances in communities which calculates the mean nearest phylogenetic neighbor among the bacterial taxa within a tick). Thus, while NTI reflects the phylogenetic structuring near the tips of the tree, NRI reflects structuring across the whole tree. The ratio between these two measures (i.e., NTI/NRI) provides a measure of phylogenetic clustering among OTUs: if NTI/NRI is positive, it suggests that there is phylogenetic clustering of OTUs (i.e., closely related OTUs are more likely to co-occur than by chance), whereas negative values indicates phylogenetic overdispersion (i.e., co-occurring OTUs are less related than expected by chance) [30]. We performed the analyses using the *picante* package [31]. To create null models, we randomized the bacterial community compositions obtained from the data by using same distance matrices, but randomizing the bacterial OTU labels across taxa.

For each alpha diversity measures we used linear mixed models with the R package *lme4* [32] to test for associations between bacterial alpha diversity and tick life stage/sex, sampling month and linear and quadratic terms of elevation with full interactions between linear terms. Sampling location was included as a random effect in the model. We used a model selection approach based on Akaike’s Information Criterion and model fit with conditional R^2^ in the package *piecewiseSEM* [33] to test which combination of factors best describes variation in tick bacterial alpha diversity. The model selection is presented in the supplementary methods (S2) while the final model is described in the results.

#### Bacterial beta taxonomic diversity

We analyzed tick bacterial beta diversity on pairwise matrices using five different indices. Two of the indices measure taxonomic beta diversity: Bray-Curtis dissimilarity, which takes into account the abundance of sequence reads [34] and Jaccard index, which takes into account only presence-absence of OTUs [35], thus providing information on both aspects of beta diversity. The other three indices measure phylogenetic beta diversity: weighted UniFrac (wUF), which takes into account the unshared branch lengths for all OTUs and weights OTUs based on OTU counts [36] equivalent to phylogenetic alpha diversity index; βNTI which is a between-community equivalent of NTI (see above) [37], and Bray-Curtis based Raup-Crick (BC-RC) abundance which measures the deviance of observed turnover while taking into account OTU relative abundances (i.e., between-community equivalent of NRI) [38]. Bray-Curtis and Jaccard indices were calculated with the R package *vegan*, UniFrac with *mothur*, βNTI with *MicEco* package [39] and BC-RC with code presented in Stegen et al. [38].

First, we performed permutational ANOVA with dissimilarity matrices using the package *vegan* to test for association between the first three measures of beta diversity (Jaccard, Bray-Curtis, Unifrac) and elevation, tick life stage/sex, sampling location and sampling month. Permutational ANOVA partitions distance matrices among sources of variation and fits linear models [40]. Initial models included all variables and interactions and, if non-significant, they were dropped during the model selection by removing the variables with the highest p-value, starting with the least significant interaction. Model selection was evaluated based on R^2^ values of remaining variables. The results of the permutational ANOVA are visualised by performing non-metrical multidimensional scaling on Bray-Curtis dissimilarities and plotting the samples on the two first axes.

Second, we used an analysis of multivariate homogeneity of group dispersion using the package *vegan* to test whether elevation, sampling location, sampling month or tick life stage/sex is associated with tick microbiota composition, again using the first three measures of beta diversity (Jaccard, Bray-Curtis, Unifrac). This analysis is a multivariate analogue of Levene’s test for homogeneity of variances [41] and tests whether variation in community composition among groups is similar.

#### Influences of ecological processes on community turnover

We used two measures of beta diversity (βNTI and BC-RC) for the analysis of ecological processes on community turnover. To study which ecological processes shape within-tick bacterial community composition, we used the phylogenetic signal of organismal niches as described by Stegen et al. [38, 42]. By assuming that closely related taxa are ecologically more similar to each other and thus their niches are more similar, we can infer which processes govern community composition. Stochastic dynamics should lead to random community assembly, environmental filtering should lead to a community consisting of taxa that are more closely related than expected by chance whereas strong competition should lead to a community consisting of less closely related taxa. Finally, environmental change should lead to increased phylogenetic turnover.

Two cases of deterministic processes are possible: if βNTI < -2, phylogenetic turnover is lower than expected by chance suggesting consistent selective pressures (*homogenous selection*), if βNTI > 2, phylogenetic turnover is higher than expected by chance, suggesting shifts in selective pressure due to environmental change (*variable selection*). If βNTI is between -2 and 2, it suggests stochastic processes determine community composition [42]. If BC-RC < -0.95, the compositional turnover between communities is low, and thus suggesting a strong dispersal between two communities (*homogenizing dispersal*). If BC-RC > 0.95, turnover is high due to a low rate of dispersal leading to ecological drift (*dispersal limitation*). Finally, in situations of moderate dispersal and weak selection, it is possible that none of these four processes shape community composition (*undominated*) [42]. We analyzed processes within sites and compared then within-site results for tick sex/stages and elevations. Due to low sample sizes per site, nymphs and samples from high elevations were not included in this analyses.

Additional analyses of phylosymbiosis between tick population genetic structure and microbiota structure and random forest classification are presented in the Supplementary material (S3 and S4).

## Results

We sequenced the microbiota of 92 *Ixodes ricinus* ticks and five negative controls, resulting in 13 214 477 amplicons. No amplification was observed in the negative controls. After contig assembly and quality control, 1 802 719 sequences were retained for OTU analysis. There was a median of 1 661 quality-controlled amplicons per tick, with an interquartile range of 5 744. 79 samples with more than 500 amplicons per sample and a Good’s coverage estimator ≥0.95 were included in the diversity analyses (Fig. 2).

**Figure 2:**
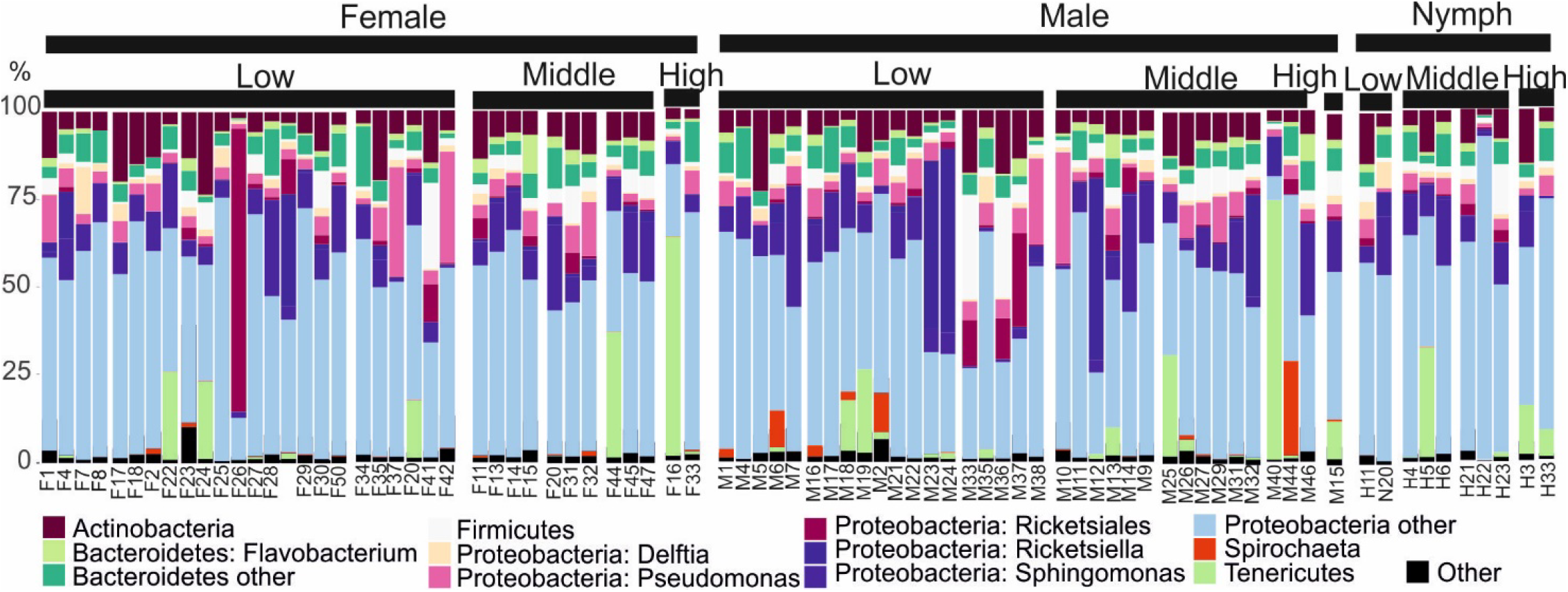
The most common bacterial taxa observed in ticks, ordered by tick sex/life stage and sampling elevations.

In total, 5 181 bacterial OTUs were identified. The median number of OTUs per rarified sample was 83 OTUs, with a 95% confidence interval of 30 - 121 OTUs. After excluding OTUs that occurred in only one sample, 864 OTUs were used in subsequent analyses. Four OTUs were present in at least 90% of the samples: *Ca*. Midichloria (Otu0001), *Pseudomonas* (Otu0002) and *Sphingomonas* (Otu0006 and Otu0009). Together, they represented 38.9 % of all amplicons. *Ca*. Midichloria was present in all samples.

### Ixodes ricinus *microbiota alpha diversity*

There was a significant elevation and sampling month effect on tick microbiota alpha diversity based on Faith’s phylogenetic index. Lower bacterial diversity was observed at higher elevations (2.0 index points per 1 000m) and diversity increased from June to August (1.74 indexpoints). For other alpha diversity indices (inverse Simpson, NRI and NTI) only tick sex / life stage was retained in the final models (Table 2, S1). Female ticks had the lowest bacterial diversity while male ticks or tick nymphs had the highest (Table 2, Fig. 3). No other environmental variables were significantly associated with tick microbiota alpha diversity (Table S1).

**Table 2:** The final linear mixed models for different alpha diversity measures.

| Measure | Variables | Value | Standard error | DF | t-value | p |
| --- | --- | --- | --- | --- | --- | --- |
| Inverse Simpson | Female intercept | 7.58 | 1.27 | 74 | 5.97 | < 0.001 |
|  | Male | 2.81 | 1.45 | 74 | 2.21 | 0.03 |
|  | Nymph | 3.41 | 2.17 | 74 | 2.11 | 0.04 |
| Faith's phylogenetic diversity | June 0 meter intercept | 1.55 | 2.64 | 74 | 0.59 | 0.56 |
|  | Elevation | -0.002 | 0.0001 | 74 | -2.02 | 0.05 |
|  | Month | 0.86 | 0.37 | 74 | 2.36 | 0.02 |
| NRI | Female intercept | -1.95 | 0.19 | 74 | -10.3 | < 0.001 |
|  | Male tick | 0.95 | 0.28 | 74 | 3.42 | 0.001 |
|  | Tick nymph | 0.88 | 0.41 | 74 | 2.15 | 0.04 |
| NTI | Female intercept | -2.17 | 0.18 | 74 | -12.2 | < 0.001 |
|  | Male tick | 0.59 | 0.26 | 74 | 2.27 | 0.02 |
|  | Tick nymph | 0.11 | 0.39 | 74 | 0.29 | 0.78 |

**Figure 3:**
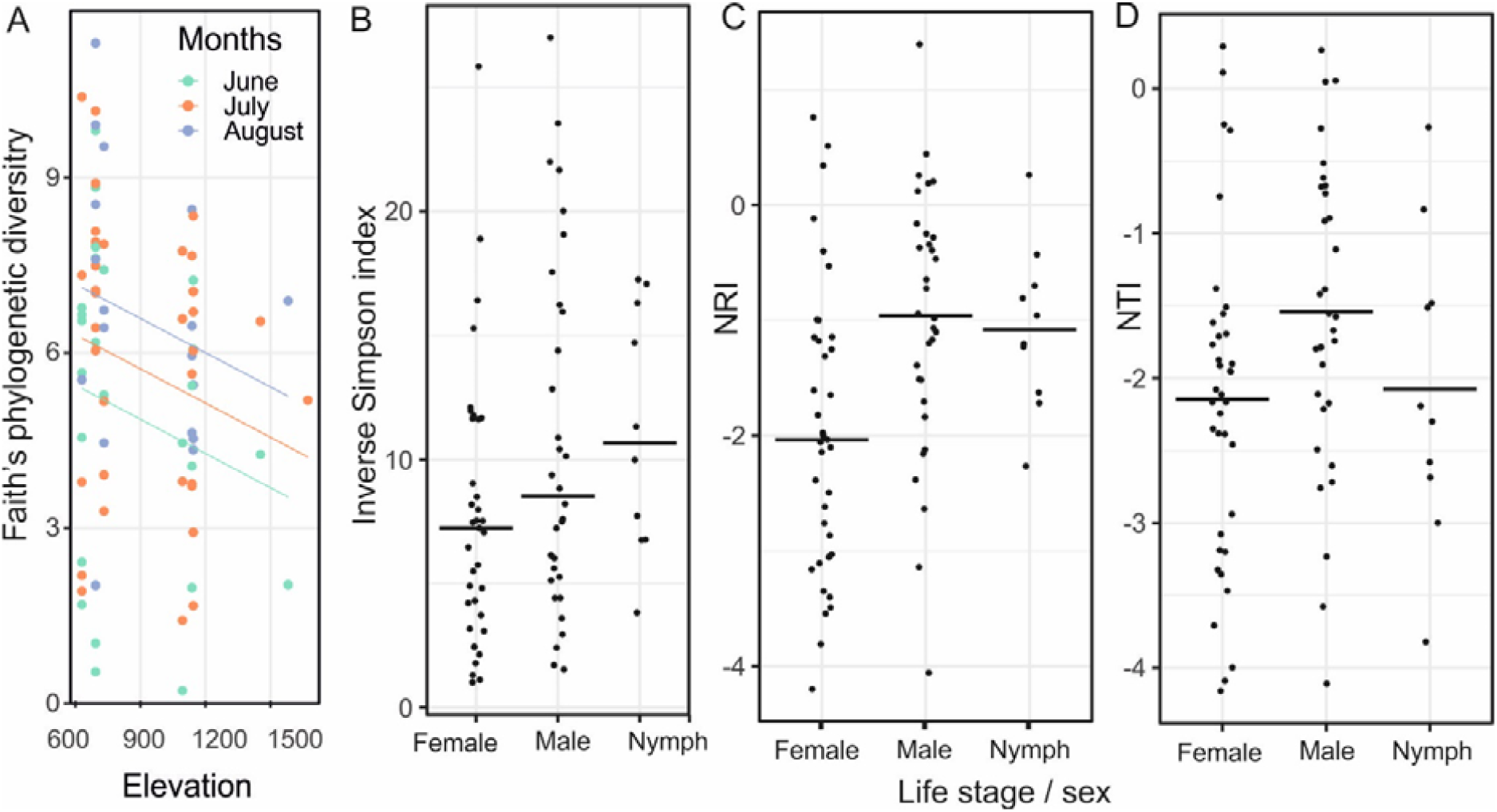
Differences in tick bacterial alpha diversity (a) as Faith’s phylogenetic diversity along elevational clines across sampling months (green: June, orange: July, purple: August) and between tick life stages / sexes when measured as (b) inverse Simpson index, (c) NRI and (d) NTI.

NTI/NRI was mostly (70/79) positive across the samples (median 1.13 with interquartile range 0.81-1.74), suggesting phylogenetic structuring of tick bacterial microbiota. There were no clear effects of elevation on phylogenetic structuring (high elevations 80%, median 1.60, IQR 0.61-2.34; middle elevations 86%, median 1.05, IQR 0.85-2.31 and low elevations 91%, median 1.19, IQR 0.77-1.65; F_5,73_=0.73, p = 0.60).

### Ixodes ricinus *microbiota beta diversity*

First, the analysis of tick microbiota beta diversity based on Jaccard index revealed significant differences in microbiota compositions along elevational clines. In addition, tick stage/sex was a significant predictor of beta diversity across all beta diversity indices (Table 3). No other variable was significantly associated with microbial beta diversity (Table S2).

**Table 3:**
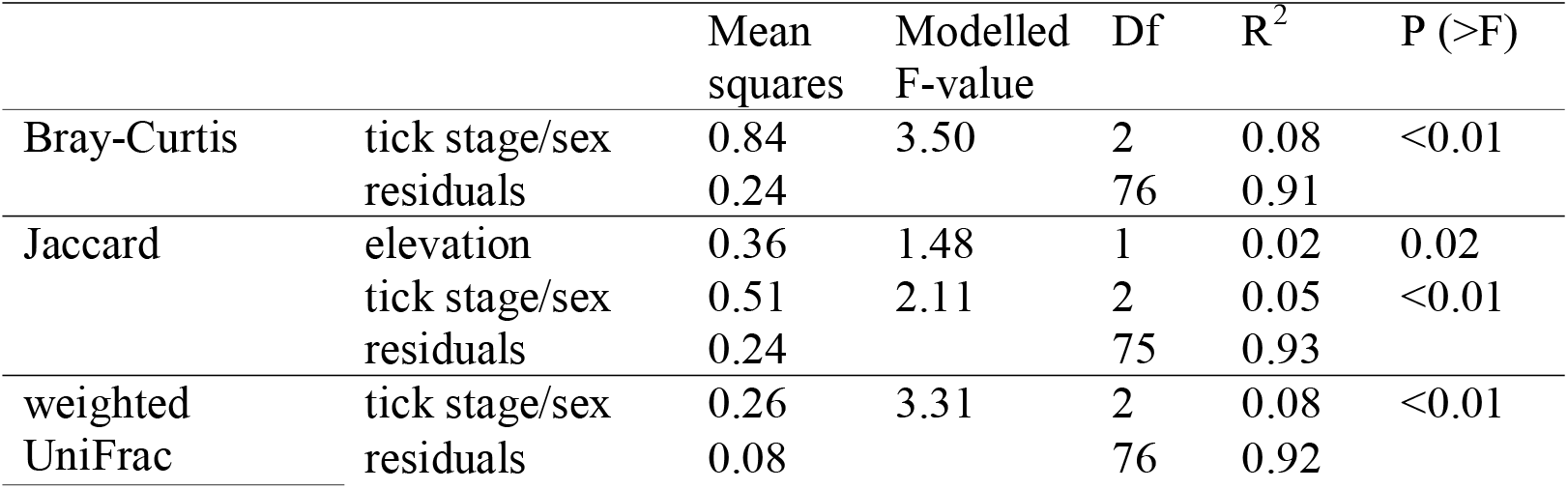
Best models describing bacterial microbiota beta diversity (quantified as Bray-Curtis dissimilarity, Jaccard distance and unweighted UniFrac distance, respectively).

Second, a significantly larger group dispersion was observed at lower elevations (Bray-Curtis dissimilarity: F_8,70_ = 5.9, adj. p < 0.001, Jaccard distance: F_8,70_ = 11.4, adj. p < 0.001; wUF: F_8,70_ = 3.1, adj. p = 0.02, Fig. 4) suggesting that among-tick variation in bacterial community composition is higher at lower elevation. Significant group dispersion was observed across sampling locations (Bray-Curtis index F_2,76_ = 5.3, adj. p = 0.02), while for other indices and variables, no significant heterogeneity in bacterial community composition was observed (Table S3).

**Figure 4:**
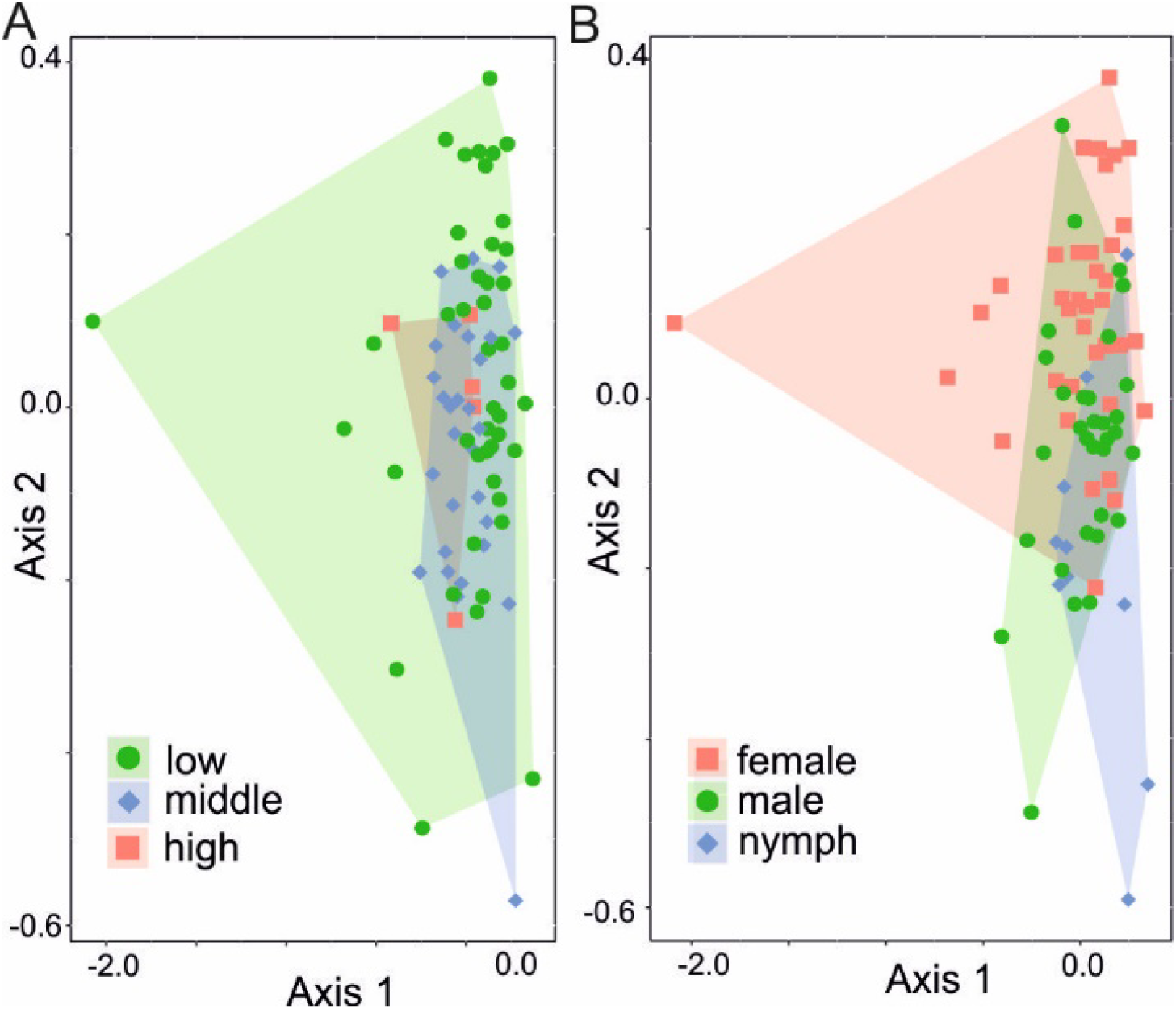
Tick microbial beta diversity along elevation clines and among tick life stages / sex and. Results are visualized by performing non-metrical multidimensional scaling on community composition and plotting the samples on the first two axes. The effect on (a) elevation and (b) tick life stage/sex on group composition can be seen by differences in areas covered by polygons. The decreasing group dispersion along elevational gradient is shown by smaller polygons in higher elevations, meaning variation in community composition is higher at lower elevations.

#### Influences of ecological processes on community turnover

In general, *Undominated* ecological processes were the most common relationships among communities (Fig. 5, Table S5-6). It suggests a moderate rate of dispersal among communities and relatively weak selection. There were no statistically significant differences in community turnover process between in samples from the same vs. different sites (χ^2^_4_ = 2.45, p = 0.65), between low and medium elevation sites (χ^2^_4_= 4.10, df = 4, p = 0.39) and between females and males (χ^2^_4_ = 6.88, p = 0.14). As between-site and within-site processes had similar distributions, the scale of the processes affecting tick microbiota is likely either larger (i.e., geographical) or smaller (i.e., within-tick) than our study. Among the groups with larger sample sizes, it is notable that female ticks showed a very low proportion of *variable selection* (1.1%), whereas *variable selection* was of substantially higher importance in male ticks (6.3%). In contrast, *homogenous selection* was more pronounced in female ticks (7.3%) compared to male ticks (3.3%). The sum of deterministic processes (i.e. *homogenous* and *variable selection* combined) was similar in females and males (8.4 % and 9.5%, respectively). It suggests that deterministic processes have a similar, though limited, effect in shaping the microbiota composition of female and male ticks, while the specific type of selection (*homogenous* vs. *variable*) differs between sexes. Stochastic processes (i.e., *dispersal limitation, homogenizing dispersal* and especially *undominated* processes) were found to shape tick bacterial community composition in all tick stages / sexes (females, 84.8%, males, 82.8% and nymphs 87.2%).

**Figure 5:**
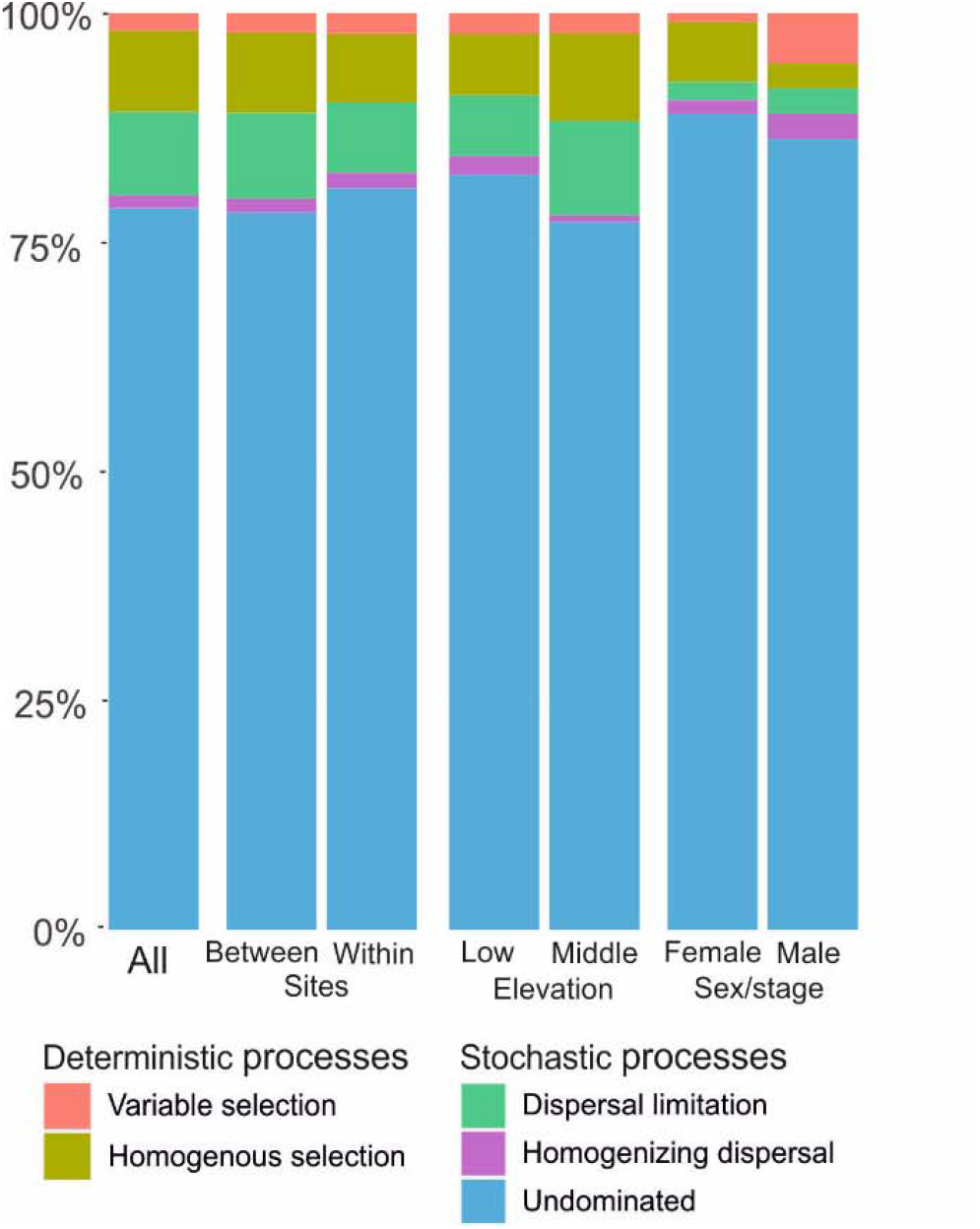
Effects of ecological processes on tick bacterial community composition across all samples, in samples between and within sites and at different elevations and across tick life stages/sexes within-sites as percentage of all pairwise comparisons. High elevations and nymphs were excluded from within-site comparisons due to low sample size.

## Discussion

We observed significant changes in bacterial alpha and beta diversity in *Ixodes ricinus* ticks along replicated elevational gradients in the Swiss Alps. Alpha diversity measured as Faith’s phylogenetic distance significantly decreased with increasing elevation, mirroring elevational diversity patterns observed in many macroscopic taxa [3] as well as in human skin microbiota [12]. In addition, microbiota composition (beta diversity) measured as Jaccard’s index differed along elevational clines with significantly lower variation in bacterial community composition at higher elevations. These differences in tick bacterial beta diversity along elevational clines contrast patterns observed in previous studies that found either no association with elevation (e.g., in soil bacteria [6]) or higher beta diversity at higher elevations (e.g., in mammalian skin and gut microbiota [11, 12]). We also observed seasonal changes in tick microbiota composition, with higher diversity later in the season.

Elevational changes in diversity were observed for some of the diversity indices, but not for others. Faith’s phylogenetic diversity measures the sum of the branch lengths of a phylogenetic tree. In contrast to taxonomic indices of alpha diversity (e.g., inverse Simpson’s index), it does not consider OTU abundances. Similarly, Jaccard beta diversity index is calculated on only presence and absence of OTU, whereas Bray-Curtis index considers OTU abundances. Thus, our results suggests that the elevational differences in tick bacterial diversity are mainly driven by the presence or absence of rarer species, rather than differences in relative OTU abundances. 40% of all amplicons belonged to only four OTUs (864 OTUs included in the analysis in total), suggesting a highly skewed abundance distribution. There is currently not a good general understanding of the functional importance of tick bacterial microbiota or how the relative abundance of different OTUs affect its functionality. While we expect that the most abundant OTUs are also the most functionally relevant, relatively rare OTUs may have substantial effects on the hosts or within-host microbial interactions [43].

Currently we can only speculate about the factors that mediate the changes in tick bacterial microbiota diversity along elevational clines. Elevation is strongly associated with temperature, soil moisture, tick host community structure and land use [1], which might all directly or indirectly shape microbial colonization and thus microbiota diversity. A recent study on *Ixodes scapularis* ticks in Canada found that ticks at the range expansion front had a different microbiota compared to ticks in the core range [44]. There is evidence that ticks have been undergoing a range expansion to higher elevations in recent years because of climate warming [15], so mountain tops represent a range expansion front [15]. Genetic diversity is typically reduced at range expansion fronts [45], which might contribute to differences in microbiota composition [10]. We directly tested for such effects in our study but found no indication that host genetic diversity or differentiation explains variation in microbial diversity (see Supplementary material S4, Table S4).

The analysis of ecological processes on community turnover suggested that stochastic processes have a strong effect on tick microbiota composition. Homogenizing dispersal, which could be facilitated by cofeeding, (i.e., two or several ticks feeding in close proximity on the same host and share microbes through their host) however, played a minor role. Ticks generally only feed from one host individual per life stage reference, which could provide opportunities for deterministic processes, yet these were rarely observed. Understanding the timeframe which shapes within-host microbiota composition will be essential to better understand the factors that contribute to the elevational differences in microbial diversity observed in our study. However, unfortunately it is next to impossible to longitudinally follow changes in tick microbiota, making this a challenging task.

In addition to elevational and seasonal effects on the bacterial microbiota of ticks we observed substantial differences in tick microbiota composition across tick life stages and sexes. In line with our findings a lower microbiota diversity in female ticks has been previously observed in *Ixodes scapularis* and *I. affinis* ticks [46, 47]. Sex differences in microbiota composition could be due to sex-differences in physiology or behaviour (including host preference). In many species males have larger home ranges than females, which might lead to exposure to a more diverse bacterial community and explain the higher microbial diversity in male ticks. Yet, to our knowledge movement patterns of male and female ticks have not been studied to date. No difference in seasonal activity is observed between adult male and female ticks [48] but during the nymph stage, female ticks become more engorged, i.e., take up more blood [49], which might partly explain the observed sex differences in microbiota composition [23].

In conclusion, we found that tick bacterial alpha diversity decreased with increasing elevation and that variation in bacterial communities is much more pronounced at lower elevations. Both of these effects were mainly driven by the presence-absence of rarer species rather than differences in relative OTU abundances. Given that bacterial microbiota composition influences the vector competence of ticks [50], understanding the functional consequences of the observed elevational differences in microbiota composition for tick-borne disease dynamics will be an important next step.

## Acknowledgements

This study was funded by Finnish Cultural Foundation Postdoc Pool grant (to TA), the Stiftung für wissenschaftliche Forschung an der Universität Zürich (17_027), the Swiss National Science Foundation (PP00P3_128386 and PP00P3_157455), the University of Zurich Research Priority Program “Evolution in Action: from Genomes to Ecosystems”, the Faculty of Science of the University of Zurich, and the Baugarten Stiftung (all to BT). We thank the numerous people who helped collecting ticks in the field, Glauco Camenisch, Elisa Granato, Jennifer Morger and Alessia Pennachia for help with laboratory work, Lucy Poveda for help with MiSeq sequencing and Frédéric Guillaume for providing access to IT infrastructure.

## References

1. Sundqvist MK, Sanders NJ, Wardle DA (2013) Community and ecosystem responses to elevational gradients: Processes, mechanisms, and insights for global change. Annu Rev Ecol Evol Syst 44:261–280. https://doi.org/10.1146/annurev-ecolsys-110512-135750

2. Körner C (2007) The use of “altitude” in ecological research. Trends Ecol Evol 22:569–574. https://doi.org/10.1016/j.tree.2007.09.006

3. Rahbek C (2005) The role of spatial scale and the perception of large-scale species-richness patterns. Ecol Lett 8:224–239. https://doi.org/10.1111/j.1461-0248.2004.00701.x

4. Dunn RR, Mccain CM, Sanders NJ (2007) When does diversity fit null model predictions? Scale and range size mediate the mid-domain effect. Glob Ecol Biogeogr 16:305–312. https://doi.org/10.1111/j.1466-8238.2006.00284.x

5. Nogués-Bravo D, Araújo MB, Romdal T, Rahbek C (2008) Scale effects and human impact on the elevational species richness gradients. Nature 453:216–219. https://doi.org/10.1038/nature06812

6. Fierer N, Mccain CM, Meir P, et al (2011) Microbes do not follow the elevational diversity patterns of plants and animals. Ecology 92:797–804. https://doi.org/10.1890/10-1170.1

7. Singh D, Lee-Cruz L, Kim WS, et al (2014) Strong elevational trends in soil bacterial community composition on Mt. Halla, South Korea. Soil Biol Biochem 68:140–149. https://doi.org/10.1016/j.soilbio.2013.09.027

8. Wang J, Soininen J, Zhang Y, et al (2011) Contrasting patterns in elevational diversity between microorganisms and macroorganisms. J Biogeogr 38:595–603. https://doi.org/10.1111/j.1365-2699.2010.02423.x

9. Looby CI, Martin PH (2020) Diversity and function fo soil microbes on montane gradients: the state of knowledge in a changing world. FEMS Microbiol Ecol 96:fiaa122 https://doi.org/10.1093/femsec/fiaa122

10. Benson AK, Kelly SA, Legge R, et al (2010) Individuality in gut microbiota composition is a complex polygenic trait shaped by multiple environmental and host genetic factors. Proc Natl Acad Sci U S A 107:18933–18938. https://doi.org/10.1073/pnas.1007028107

11. Li H, Zhou R, Zhu J, et al (2019) Environmental filtering increases with elevation for the assembly of gut microbiota in wild pikas. Microb Biotechnol 12:976–992. https://doi.org/10.1111/1751-7915.13450

12. Li H, Wang Y, Yu Q, et al (2019) Elevation is associated with human skin microbiomes. Microorganisms 7:. https://doi.org/10.3390/microorganisms7120611

13. Ross BD, Hayes B, Radey MC, et al (2018) Ixodes scapularis does not harbor a stable midgut microbiome. ISME J 12:2596–2607. https://doi.org/10.1038/s41396-018-0161-6

14. Narasimhan S, Rajeevan N, Liu L, et al (2014) Gut microbiota of the tick vector Ixodes scapularis modulate colonization of the Lyme disease spirochete. Cell Host Microbe 15:58–71. https://doi.org/10.1016/j.chom.2013.12.001

15. Garcia-Vozmediano A, Krawczyk AI, Sprong H, et al (2020) Ticks climb the mountains: Ixodid tick infestation and infection by tick-borne pathogens in the Western Alps. Ticks Tick Borne Dis 11:101489. https://doi.org/10.1016/j.ttbdis.2020.101489

16. Mysterud A, Jore S, Østerås O, Viljugrein H (2017) Emergence of tick-borne diseases at northern latitudes in Europe: a comparative approach. Sci Rep 7:16316. https://doi.org/10.1038/s41598-017-15742-6

17. Gray JS (1991) The development and seasonal activity of the tick Ixodes ricinus: a vector of Lyme borreliosis. Rev Med Vet Entomol 79:323–333

18. Gardiner WP, Gettinby G, Gray JS (1981) Models based on weather for the development phases of the sheep tick, Ixodes ricinus L. Vet Parasitol 9:75–86. https://doi.org/10.1016/0304-4017(81)90009-1

19. Oechslin CP, Heutschi D, Lenz N, et al (2017) Prevalence of tick-borne pathogens in questing Ixodes ricinus ticks in urban and suburban areas of Switzerland. Parasites and Vectors 10:1–18. https://doi.org/10.1186/s13071-017-2500-2

20. Estrada-Peña A, Ostfeld RS, Peterson a T, et al (2014) Effects of environmental change on zoonotic disease risk: an ecological primer. Trends Parasitol 30:205–14. https://doi.org/10.1016/j.pt.2014.02.003

21. Aivelo T, Norberg A, Tschirren B (2019) Bacterial microbiota composition of Ixodes ricinus ticks: the role of environmental variation, tick characteristics and microbial interactions. PeerJ 7:e8217. https://doi.org/10.1101/559245

22. Thapa S, Zhang Y, Allen MS (2019) Effects of temperature on bacterial microbiome composition in Ixodes scapularis ticks. Microbiologyopen 8:e719. https://doi.org/10.1002/mbo3.719

23. Narasimhan S, Swei A, Abouneameh S, et al (2021) Grappling with the tick microbiome. Trends Parasitol. https://doi.org/10.1016/j.pt.2021.04.004

24. Lemoine M, Cornetti L, Tschirren B (2018) Does Borrelia burgdorferi sensu lato facilitate the colonisation of marginal habitats by Ixodes ricinus? A correlative study in the Swiss Alps. bioRxiv. https://doi.org/10.1101/273490

25. Carey H V, Walters WA, Knight R (2013) Seasonal restructuring of the ground squirrel gut microbiota over the annual hibernation cycle. Am J Physiol Regul Integr Comp Physiol 304:R33–42. https://doi.org/10.1152/ajpregu.00387.2012

26. Kozich JJ, Westcott SL, Baxter NT, et al (2013) Development of a dual-index sequencing strategy and curation pipeline for analyzing amplicon sequence data on the MiSeq Illumina sequencing platform. Appl Environ Microbiol 79:5112–20. https://doi.org/10.1128/AEM.01043-13

27. Simpson EH (1949) Measurement of diversity. Nature 163:688

28. Faith DP (1992) Conservation evaluation and phylogenetic diversity. Biol Conserv 61:1–10. https://doi.org/10.1016/0003-2697(75)90168-2

29. Oksanen J, Blanchet FG, Kindt R, et al (2020) vegan: Community Ecology Package. Version 2.5-7.

30. Webb CO (2000) Exploring the phylogenetic structure of ecological communities: An example for rain forest trees. Am Nat 156:145–155. https://doi.org/10.1086/303378

31. Kembel SW, Cowan PD, Helmus MR, et al (2010) Picante: R tools for integrating phylogenies and ecology. Bioinformatics 26:1463–1464. https://doi.org/10.1093/bioinformatics/btq166

32. Bates D, Mächler M, Bolker B, Walker S (2015) Fitting linear mixed-effects models using lme4. J Stat Softw 67:p. https://doi.org/10.18637/jss.v067.i01

33. Lefcheck JS (2016) piecewiseSEM: Piecewise structural equation modelling in R for ecology, evolution, and systematics. Methods Ecol Evol 7:573–579. https://doi.org/10.1111/2041-210X.12512

34. Bray JR, Curtis JT (1957) An ordination of the upland forest community of Southern Wisconsin. Ecol Monogr 27:325–349. https://doi.org/10.2307/1942268

35. Jaccard P (1912) The distribution of the flora in the alpine zone. New Phytol 11:37–50. https://doi.org/10.1111/j.1469-8137.1912.tb05611.x

36. Lozupone C, Lladser ME, Knights D, et al (2011) UniFrac: An effective distance metric for microbial community comparison. ISME J 5:169–172. https://doi.org/10.1038/ismej.2010.133

37. Chase JM, Kraft NJB, Smith KG, et al (2011) Using null models to disentangle variation in community dissimilarity from variation in α-diversity. Ecosphere 2:p. https://doi.org/10.1890/ES10-00117.1

38. Stegen JC, Lin X, Fredrickson JK, et al (2013) Quantifying community assembly processes and identifying features that impose them. ISME J 7:2069–2079. https://doi.org/10.1038/ismej.2013.93

39. Russel J. (2021). Russel88/MicEco (Version v0.9.14). Zenodo. http://doi.org/10.5281/zenodo.4639787

40. Anderson MJ (2001) A new method for non-parametric multivariate analysis of variance. Austral Ecol 26:32–46. https://doi.org/10.1046/j.1442-9993.2001.01070.x

41. Anderson MJ, Ellingsen KE, McArdle BH (2006) Multivariate dispersion as a measure of beta diversity. Ecol Lett 9:683–693. https://doi.org/10.1111/j.1461-0248.2006.00926.x

42. Stegen JC, Lin X, Fredrickson JK, Konopka AE (2015) Estimating and mapping ecological processes influencing microbial community assembly. Front Microbiol 6:1–15. https://doi.org/10.3389/fmicb.2015.00370

43. Lozupone C a, Stombaugh JI, Gordon JI, et al (2012) Diversity, stability and resilience of the human gut microbiota. Nature 489:220–30. https://doi.org/10.1038/nature11550

44. Clow KM, Weese JS, Rousseau J, Jardine CM (2017) Microbiota of field-collected Ixodes scapularis and Dermacentor variabilis from eastern and southern Ontario, Canada. Ticks Tick Borne Dis 9:235–244. https://doi.org/10.1016/j.ttbdis.2017.09.009

45. Excoffier L, Foll M, Petit RJ (2009) Genetic consequences of range expansions. Annu Rev Ecol Evol Syst 40:481–501. https://doi.org/10.1146/annurev.ecolsys.39.110707.173414

46. van Treuren W, Ponnusamy L, Brinkerhoff RJ, et al (2015) Variation in the microbiota of Ixodes ticks with regard to geography, species, and sex. Appl Environ Microbiol 81:6200–6209. https://doi.org/10.1128/AEM.01562-15

47. Zolnik CP, Prill RJ, Falco RC, et al (2016) Microbiome changes through ontogeny of a tick pathogen vector. Mol Ecol 25:4963–4977. https://doi.org/10.1111/mec.13832

48. Randolph S, Green R, Hoodless A, Peacey M (2002) An empirical quantitative framework for the seasonal dynamics of Ixodes ricinus. Int J Paras 32:979–989

49. Dusbábek F (1996) Nymphal sexual dimorphism in the sheep tick Ixodes ricinus (Acari: Ixodidae). Folia Parasitol 43:75–79

50. Narasimhan S, Fikrig E (2015) Tick microbiome: the force within. Trends Parasitol 31:315–323. https://doi.org/10.1016/j.pt.2015.03.010

