## Supplementary Material for "Elevational changes in bacterial microbiota structure and diversity in an arthropod-disease vector"

Tuomas Aivelo, Mélissa Lemoine & Barbara Tschirren

### *S1: Supplementary data collection*

Tick microbiota sequencing

16S sequencing libraries were prepared following the Earth Microbiome 16S Illumina Amplicon protocol, using the primers

515FB: CTTTCCCTACACGACGCTCTTCCGATCTNNNNNNNNGTGYCAGCMGCCGCGGTAA
806RB: GGAGTTCAGACGTGTGCTCTTCCGATCTNNNNNNNNGGACTACNVGGGTWTCTAAT

which also contained the Illumina Miseq primer binding site [1–3]. Samples and negative controls were randomized across two plates. Each reaction contained 1 µl of template DNA with 0.5 µM forward and reverse primers, 0.3 mM dNTP, 0.4 µl KAPA HiFi HotStart polymerase (KAPA Biosystems, Basel, Switzerland) in a final volume of 20 µl. The PCR protocol started with an initial denaturation at 95 C° for 3 minutes, followed by 35 cycles of 20 seconds at 98 C°, 60 seconds at 50 C° and 30 seconds at 72 C° with final elongation step of 5 minutes at 72 C°. We ran PCR products on 2% agarose gels, cut out the bands corresponding to the amplicon size (350 bp) and eluted them in 10 µl milli-Q water. We then purified the extracted amplicons using MinElute Gel Extraction kit (Qiagen, Hilden, Germany) following the manufacturer's protocol. In second PCR step Illumina Miseq adaptors and dual indexing Miseq barcodes were incorporated into the previously amplified V3-V4 regions. The second PCR protocol started with an initial denaturation at 95 C° for 3 minutes, followed by 15 cycles of 20 seconds at 98 C°, 60 seconds at 54 C° and 30 seconds at 72 C° with final elongation step of 5 minutes at 72 C°. Products were visualized and purified as described above. We then measured the DNA concentration of the purified product with Invitrogen Qubit 4 Fluorometer ssDNA assay kit (Thermo Fisher Scientific, Walthan, MA, USA) and created library pools by mixing equimolar amounts of purified products to reach a 4 nM library concentration. The libraries were further processed using standard normalization protocols with PhiX controls and sequenced on a Illumina MiSeq at the Functional Genomic Center Zurich using reagent kit 600cycle v3 (Illumina, San Diego, CA, USA) with a target length of 250 bp per amplicon following the manufacturer's protocol.

### *S2: Supplementary methods and results of diversity analysis*

Alpha diversity

**Table S1a**: Model selection for modelling alpha diversity of tick bacterial microbiota (inverse Simpson index).

<table>
<tr><th>Model</th><th>AIC</th><th>cond. $R^2$</th></tr>
<tr><td>invsimpson ~ masl*tick_stage*month+masl^2, random= ~1|location</td><td>584.0</td><td>0.20</td></tr>
<tr><td>invsimpson ~ masl*tick_stage*month, random= ~1|location</td><td>561.4</td><td>0.20</td></tr>
<tr><td>invsimpson ~ masl+month+tick_stage+masl:month+tick_stage:month+ masl:tick_stage, random= ~1|location</td><td>547.0</td><td>0.15</td></tr>
<tr><td>invsimpson ~ masl+month+tick_stage+masl:tick_stage+tick_stage:month, random= ~1|location</td><td>527.4</td><td>0.15</td></tr>
<tr><td>invsimpson ~ masl+month+tick_stage++tick_stage:month, random= ~1|location</td><td>530.9</td><td>0.15</td></tr>
<tr><td>invsimpson ~ masl+tick_stage+tick_stage:month , random= ~1|location</td><td>516.3</td><td>0.15</td></tr>
<tr><td>invsimpson ~ masl+tick_stage, random= ~1|location</td><td>522.9</td><td>0.13</td></tr>
<tr><td>invsimpson ~ tick_stage, random= ~1|location</td><td>511.9</td><td>0.12</td></tr>
<tr><td>invsimpson ~ 1, random= ~1|location</td><td>515.9</td><td>0.05</td></tr>
</table>

**Table S1b**: Model selection for modelling alpha diversity of tick bacterial microbiota (Faith's index).

<table>
<tr><th>Model</th><th>AIC</th><th>cond. $R^2$</th></tr>
<tr><td>phylodiv ~ masl*tick_stage*month+masl^2, random= ~1|location</td><td>459.8</td><td>0.21</td></tr>
<tr><td>phylodiv ~ masl*tick_stage*month, random= ~1|location</td><td>436.2</td><td>0.21</td></tr>
<tr><td>phylodiv ~ masl+month+tick_stage+masl:month+tick_stage:month+ masl:tick_stage, random= ~1|location</td><td>417.0</td><td>0.17</td></tr>
<tr><td>phylodiv ~ masl+month+tick_stage+masl:tick_stage+tick_stage:month, random= ~1|location</td><td>395.2</td><td>0.14</td></tr>
<tr><td>phylodiv ~ masl+month+tick_stage+tick_stage:month, random= ~1|location</td><td>382.7</td><td>0.13</td></tr>
<tr><td>phylodiv ~ masl+tick_stage+ month , random= ~1|location</td><td>382.6</td><td>0.13</td></tr>
<tr><td>phylodiv ~ masl+month, random= ~1|location</td><td>381.2</td><td>0.12</td></tr>
<tr><td>phylodiv ~ masl, random= ~1|location</td><td>391.2</td><td>0.08</td></tr>
<tr><td>phylodiv ~~ 1, random= ~1|location</td><td>393.6</td><td>0.04</td></tr>
</table>

**Table S1c**: Model selection for modelling alpha diversity of tick bacterial microbiota (NRI).

<table>
<tr><th>Model</th><th>AIC</th><th>cond. $R^2$</th></tr>
<tr><td>NRI ~ masl*tick_stage*month+masl^2, random= ~1|location</td><td>362.8</td><td>0.19</td></tr>
<tr><td>NRI ~ masl+month+tick_stage+masl:month+tick_stage:month+ masl:tick_stage+ masl^2, random= ~1|location</td><td>343.3</td><td>0.12</td></tr>
<tr><td>NRI ~ masl+month+tick_stage+masl:month+tick_stage:month+ masl:tick_stage</td><td></td><td></td></tr>
<tr><td>NRI ~ masl+month+tick_stage+masl:month+ masl:tick_stage, random= ~1|location</td><td>317.2</td><td>0.13</td></tr>
<tr><td>NRI ~ masl+month+tick_stage+ masl:tick_stage, random= ~1|location</td><td>299.1</td><td>0.10</td></tr>
<tr><td>NRI ~ masl+tick_stage+ masl:tick_stage, random= ~1|location</td><td>295.5</td><td>0.19</td></tr>
<tr><td>NRI ~ tick_stage+masl, random= ~1|location</td><td>272.6</td><td>0.06</td></tr>
<tr><td>NRI ~ tick_stage, random= ~1|location</td><td>257.0</td><td>0.06</td></tr>
<tr><td>NRI ~ 1, random= ~1|location</td><td>264.9</td><td>0.00</td></tr>
</table>

**Table S1d**: Model selection for modelling alpha diversity of tick bacterial microbiota (NTI).

| Model | AIC | cond. $R^2$ |
|---|---|---|
| NTI ~ masl*tick_stage*month+ masl^2, random= ~1\|location | 352.4 | 0.14 |
| NTI ~ masl+month+tick_stage+masl:month+tick_stage:month+ masl:tick_stage+masl^2, random= ~1\|location | 327.2 | 0.14 |
| NTI ~ masl+month+tick_stage++tick_stage:month+ masl:tick_stage+ masl^2, random= ~1\|location | 312.4 | 0.14 |
| NTI ~ masl+month+tick_stage+masl:tick_stage+ masl^2, random= ~1\|location | 306.5 | 0.14 |
| NTI ~ masl+month+tick_stage+ masl^2, random= ~1\|location | 281.1 | 0.12 |
| NTI ~ masl+tick_stage+month , random= ~1\|location | 254.0 | 0.11 |
| NTI ~ month+tick_stage, random= ~1\|location | 238.7 | 0.11 |
| **NTI ~ tick_stage, random= ~1\|location** | **236.3** | **0.10** |
| NTI ~ 1, random= ~1\|location | 249.4 | 0.00 |

Beta diversity

**Table S2a**: Model selection for modelling Bray-Curtis beta diversity of tick bacterial microbiota using permutational ANOVA. The final model is highlighted in bold.

| Model | Significant variables | $R^2$ |
|---|---|---|
| betad~tick_stage*masl*location*month | tick_stage, location:month | 0.46 |
| betad~tick_stage+masl*location*month+tick_stage:masl:location:month+tick_stage:masl:month+tick_stage:location:month+tick_stage:masl+tick_stage:month+tick_stage:location | tick_stage, location:month | 0.44 |
| betad~tick_stage+masl*location*month+ tick_stage:masl:month+tick_stage:location:month+tick_stage:masl+tick_stage:month+tick_stage:location | tick_stage, location:month | 0.42 |
| betad~masl+tick_stage+location*month+tick_stage:masl:month+tick_stage:location:month+tick_stage:masl+tick_stage:month+tick_stage:location+masl:location:month | masl, location, tick_stage, masl:location | 0.39 |
| betad~masl+tick_stage+location*month+tick_stage:masl:month+tick_stage:masl+tick_stage:month+tick_stage:location+masl:location:month | tick_stage, location:month | 0.39 |
| betad~masl+tick_stage+location+location:month+tick_stage:masl:month+tick_stage:masl+tick_stage:month+tick_stage:location+masl:location:month | masl, location, tick_stage, location:month | 0.37 |
| betad~masl+tick_stage+location+location:month+tick_stage:masl:month+tick_stage:masl+tick_stage:month+tick_stage:location | masl, location, tick_stage, location:month, masl:tick_stage | 0.35 |
| betad~masl+tick_stage+location+location:month+tick_stage:masl+tick_stage:month+tick_stage:location | location, tick_stage, location:month, masl:tick_stage | 0.31 |
| betad~masl+tick_stage+location+location:month+tick_stage:masl+tick_stage:month | tick_stage, location:month | 0.25 |
| betad~ masl+location+tick_stage+location:month+tick_stage:month | location, tick_stage, location:month | 0.21 |
| betad~masl+tick_stage+location+location:month | tick_stage | 0.19 |
| betad~masl+tick_stage+location | tick_stage | 0.13 |
| betad~ tick_stage+location | tick_stage | 0.11 |
| **betad~ tick_stage** | **tick_stage** | **0.09** |

**Table S2b**: Model selection for modelling Jaccard index beta diversity of tick bacterial microbiota using permutational ANOVA. The final model is highlighted in bold.

| Model | Significant variables | $R^2$ |
|---|---|---|
| betad~tick_stage*masl*location*month | tick_stage, masl, tick_stage:location | 0.41 |
| betad~tick_stage+masl*location*month+tick_stage:masl:location:month+tick_stage:masl:month+tick_stage:location:month+tick_stage:masl+tick_stage:month+tick_stage:location | tick_stage, masl, tick_stage:location | 0.37 |
| betad~tick_stage+masl*location*month+ tick_stage:masl:month+tick_stage:location:month+tick_stage:masl+tick_stage:month+tick_stage:location | tick_stage, masl, tick_stage:masl:month | 0.33 |
| betad~tick_stage+ location+masl+location:month+tick_stage:masl:month+tick_stage:location:month+tick_stage:masl+tick_stage:month+tick_stage:location+masl:location:month+masl:location | tick_stage, masl, tick_stage:location | 0.33 |
| betad~tick_stage+ location+masl+location:month+tick_stage:masl:month+tick_stage:masl+tick_stage:month+tick_stage:location+masl:location:month +masl:location | tick_stage, masl tick_stage:location, tick_stage:masl:month | 0.31 |
| betad~tick_stage+ location+masl+location:month+tick_stage:masl:month+tick_stage:masl+tick_stage:month+tick_stage:location+masl:location | tick_stage, masl, tick_stage:location | 0.28 |
| betad~tick_stage+ location+masl+tick_stage:masl:month+tick_stage:masl+tick_stage:month+tick_stage:location+masl:location | tick_stage, masl | 0.25 |
| betad~tick_stage+ location+masl+tick_stage:masl+tick_stage:month+tick_stage:location+masl:location | tick_stage, masl | 0.21 |
| betad~tick_stage+location+masl+tick_stage:masl+ tick_stage:location+masl:location | tick_stage, masl | 0.17 |
| betad~tick_stage+location+ masl+ tick_stage:masl | tick_stage, masl | 0.11 |
| betad~tick_stage+masl+ masl:tick_stage | tick_stage, masl | 0.09 |
| **betad~tick_stage+masl** | **tick_stage, masl** | **0.07** |

**Table S2c**: Model selection for modelling weighted UniFrac index beta diversity of tick bacterial microbiota using permutational ANOVA. The final model is highlighted in bold.

| Model | Significant variables | $R^2$ |
|---|---|---|
| betad~tick_stage*masl*location*month | tick_stage, location:month | 0.44 |
| betad~tick_stage+masl*location*month+tick_stage:masl:location:month+tick_stage:masl:month+tick_stage:location:month+tick_stage:masl+tick_stage:month+tick_stage:location | tick_stage, location:month | 0.40 |
| betad~tick_stage+masl*location*month+ tick_stage:masl:month+tick_stage:location:month+tick_stage:masl+tick_stage:month+tick_stage:location | tick_stage, location:month | 0.37 |
| betad~tick_stage+ location+masl+location:month+tick_stage:masl:month+tick_stage:location:month+tick_stage:masl+tick_stage:month+tick_stage:location+masl:location:month+masl:location | tick_stage, location:month | 0.35 |
| betad~tick_stage+ location+masl+location:month+tick_stage:masl:month+tick_stage:masl+tick_stage:month+tick_stage:location+masl:location:month +masl:location | tick_stage, location, location:month | 0.33 |

| | | |
|---|---|---|
| betad~tick_stage+ location+masl+location:month+tick_stage:masl:month+tick_stage:masl+tick_stage:month+tick_stage:location+masl:location | tick_stage, masl | 0.31 |
| betad~tick_stage+ location+masl+tick_stage:masl:month+tick_stage:masl+tick_stage:month+tick_stage:location+masl:location | tick_stage, location | 0.28 |
| betad~tick_stage+ location+masl+tick_stage:masl+tick_stage:month+tick_stage:location+masl:location | tick_stage, location | 0.24 |
| betad~tick_stage+location+masl+tick_stage:masl+ tick_stage:location+masl:location | tick_stage, location | 0.21 |
| betad~tick_stage+location+ masl+ tick_stage:masl | tick_stage | 0.17 |
| betad~tick_stage+masl+ masl:tick_stage | tick_stage | 0.13 |
| betad~tick_stage+masl | tick_stage | 0.11 |
| **betad~tick_stage** | **tick_stage** | **0.08** |

**Table S3**: Associations between the ecological variables and the group dispersion of tick microbiota diversity. As it can only be performed with one variable at a time, we corrected for multiple testing with Hochberg's correction [4].

| Diversity index | Variable | | Mean of squares | F-value | Df | Pr (>F) | Adjusted Pr (>F) |
|---|---|---|---|---|---|---|---|
| Bray-Curtis | **masl** | **Groups** | **0.06** | **5.9** | **8** | **< 0.001** | **<0.001** |
| | | **Residuals** | **0.01** | | **70** | | |
| | **location** | **Groups** | **0.06** | **5.3** | **2** | **0.007** | **0.02** |
| | | **Residuals** | **0.01** | | **76** | | |
| | tick_stage | Groups | 0.01 | 0.81 | 2 | 0.44 | 0.45 |
| | | Residuals | 0.01 | | 76 | | |
| | month | Groups | 0.01 | 0.93 | 2 | 0.40 | 0.45 |
| | | Residuals | 0.01 | | 76 | | |
| Jaccard | **masl** | **Groups** | **0.04** | **11.44** | **8** | **< 0.001** | **< 0.001** |
| | | **Residuals** | **0.01** | | **70** | | |
| | location | Groups | 0.01 | 3.48 | 2 | 0.04 | 0.11 |
| | | Residuals | 0.01 | | 76 | | |
| | tick_stage | Groups | 0.01 | 2.54 | 2 | 0.09 | 0.17 |
| | | Residuals | 0.01 | | 76 | | |
| | month | Groups | 0.01 | 1.67 | 2 | 0.19 | 0.19 |
| | | Residuals | 0.01 | | 76 | | |
| Weighted Unifrac | **masl** | **Groups** | **0.02** | **3.11** | **8** | **0.01** | **0.02** |
| | | Residuals | 0.01 | | 70 | | |
| | location | Groups | 0.02 | 2.68 | 2 | 0.07 | 0.22 |
| | | Residuals | 0.01 | | 76 | | |
| | tick_stage | Groups | 0.01 | 0.56 | 2 | 0.57 | 0.57 |
| | | Residuals | 0.01 | | 76 | | |
| | month | Groups | 0.01 | 1.20 | 2 | 0.31 | 0.56 |
| | | Residuals | 0.01 | | 76 | | |

### *S3: Supplementary methods and results of random forest analysis*

We used Breiman's Random Forest Classifier approach with the package *randomForest* [5] to assess how well our variables of interested delimit different categories. In other words, this test assesses whether it can be predicted from the bacterial community composition whether it originates from a tick collected in a specific month, elevation or tick life stage/sex. We ran the algorithm for each of the previously mentioned variables and measured the predictive accuracy by out-of-bag error rate [5].

The random forest analysis provided an out-of-bag score of 43.0% for elevation (classification error for low elevation 0.11, for middle elevation 0.83 and for high elevation 1.00), 34.2% for sampling month (classification error for June 0.72, for July 0.17 and for August 1.00), 27.5% for tick stage/sex (classification error for females 0.19, for males 0.41 and for nymphs 0.80) and 32.0 % for sampling location (classification error for Sagogn 0.29, for Roedels 0.55 and for Passug 1.00), which corresponds to the expected percentages based on random guessing. It suggests that neither of these variables can be reliably predicted based on OTU composition.

### *S4: Supplementary methods and results of tick population genetic analyses*

To calculate genetic distances among individual ticks, neutral genetic diversity was determined as described in Aivelo et al. [6]. In short, we genotyped ticks at 11 microsatellite markers in two multiplexed amplifications and used the package *poppr* [7] in R 3.4.1 [8] to calculate Prevosti's distance between individual ticks [9].

To test for phylosymbiosis, we analyzed whether the neutral genetic distance between ticks explains differences in their bacterial communities. The association between the genetic distances among individual ticks and genetic distances of their bacterial microbiotas was calculated in three different ways: First, we correlated the dissimilarities of tick microbiota (wUF) and genetic distances among ticks and fitted loess (local polynomial regression lines; Cleveland et al. 1992) with $\alpha = 0.2$. Second, we tested for an association between the microbiota weighted UniFrac matrix and the tick genetic distance matrix using Mantel test [11]. Third, we compared trees created based on tick neutral genetic distances and tree based on tick microbiota UniFrac distances by calculating the Matching Split Distance [12, 13] for unrooted binary tree. Matching Split Distance compares tree branches implied by a pair of splits between the two trees and calculates the number of taxa that must be moved from one branch to another to make two splits identical. UniFrac-based trees with branch tip labels randomly assigned 10,000 times were used as a null models for this analysis.

No evidence for phylosymbiosis between ticks and their bacterial microbiota community was found across tests. Pairwise neutral genetic distances between individual ticks did not correlate with the dissimilarities in within-tick microbiota composition (i.e., weighted UniFrac) (LOESS curves: pseudo-$R^2$ = 0.40), suggesting that neutral genetic differences among ticks do not explain variation in within-tick community composition. Likewise, the comparison of tick genetic distance and within-tick microbial community distance matrices did not show a significant association (Mantel test: r = -0.05, p = 0.80). Finally, the phylogenetic tree of sampled ticks and the tree based on the dissimilarities of their microbiota were not significantly similar (Matching Split Distance: required number of substitutions is 583, p = 0.32 compared to null model), meaning that differentiation in tick microbiota did not reflect the genetic differentiation in ticks.

Tick population differentiation across sites, elevations and tick stage/sex were evaluated by calculating pairwise $F_{ST}$ values according to Nei [14] with the package *heirfstat* [15]. We performed Mantel tests [11] with 9999 permutations in the package *ade4* [16] to test for isolation by distance and isolation by elevation. There was no evidence for isolation by distance or isolation by elevation as no statistically significant differences were found in Mantel tests (IBD: r = - 0.29, $p_{simulated}$ = 0.95; IBE: r = 0.02, $p_{simulated}$ = 0.44).

**Table S4a:** Pairwise $F_{ST}$ values based on tick stage/sex.

| | F | M |
|---|---|---|
| M | - 0.001 | |
| N | 0.009 | 0.008 |

**Table S4b:** Pairwise $F_{ST}$ values based on site

| | Rodels | Tomils | Feldis | Sagogn | Flims | Ruschein | Passug | Castiel |
|---|---|---|---|---|---|---|---|---|
| Tomils | 0.022 | | | | | | | |
| Feldis | 0.064 | 0.017 | | | | | | |
| Sagogn | 0.022 | -0.001 | -0.044 | | | | | |
| Flims | 0.015 | -0.001 | -0.071 | -0.017 | | | | |
| Ruschein | 0.025 | 0.020 | 0.030 | -0.008 | 0.001 | | | |
| Passug | 0.035 | 0.003 | -0.054 | 0.001 | 0.002 | -0.004 | | |
| Castiel | 0.042 | 0.238 | 0.041 | -0.001 | 0.009 | 0.036 | 0.036 | |
| Praden | 0.010 | 0.034 | 0.012 | 0.005 | -0.025 | -0.079 | 0.059 | 0.032 |

**Table S4c:** Pairwise $F_{ST}$ values based on elevation

| | High | Low |
|---|---|---|
| Low | 0.0033 | |

| | | |
|---|---|---|
| Middle | 0.0184 | - 0.028 |

Based on the neutral microsatellite markers, there was thus no obvious differentiation in tick populations due to either geographic or altitudinal distances. This suggests that the difference in microbiota across life stage/sex is not due to different dispersal or differences in population composition of nymph, males or females or along elevational gradient. Furthermore, this lack of population structure could explain why there was no evidence of phylosymbiosis between tick populations and microbiotal structure.

### *S5: Supplementary results of influences of ecological processes on community turnover*

**Table S5**: The median values and interquartile range of βNTI and Bray-Curtis-based Raup-Crick for the whole sample and divided by elevation classes and tick stage/sex.

| | Whole sample | Elevation | | | Tick stage/sex | | |
|---|---|---|---|---|---|---|---|
| | | Low | Middle | High | Female | Male | Nymph |
| βNTI | -0.29<br>*-1.20 – 0.80* | – 0.32<br>*-1.16 – 0.69* | – 0.20<br>*– 1.09 – 1.10* | 1.09<br>*0.40 – 1.30* | -0.71<br>*-1.55 – 0.35* | -0.20<br>*-1.09 – 1.10* | 0.78<br>*-0.10 – 1.15* |
| Raup-Crick | 0.51<br>*0.00 – 0.80* | 0.49<br>*0.00 – 0.77* | 0.47<br>*0.01 – 0.73* | 0.28<br>*0.01 – 0.73* | 0.19<br>*-0.34 – 0.62* | 0.47<br>*0.01 – 0.73* | 0.87<br>*0.42 – 0.97* |

**Table S6**: The number of occurrences of each type of ecological processes in between-pair comparisons in all samples, between different sites, within sites in elevation and tick stage/sex categories. The numbers are crude numbers of between-pair comparisons. Rule describes how between pair comparisons are assigned to one of the five processes. Samples from high elevations and nymphs were excluded from between and within-sites comparisons due to low sample size per category, but they were included in 'All'.

| Process | | Rule | All | Between-site | Within-site | | | | |
|---|---|---|---|---|---|---|---|---|---|
| | | | | Total | Total | By elevation | | By tick stage/sex | |
| Type | Name | | | | | Low | Middle | Female | Male |
| Deterministic | | | | | | | | | |
| | Variable selection | βNTI > 2 | 59 | 38 | 11 | 8 | 3 | 2 | 4 |
| | Homogenous selection | βNTI < -2 | 273 | 157 | 36 | 23 | 13 | 13 | 2 |
| Stochastic | | | | | | | | | |
| | Homogenizing dispersal | -2 < βNTI < 2 & BC-RC < -0.95 | 44 | 27 | 9 | 7 | 1 | 4 | 2 |
| | Dispersal limitation | -2 < βNTI < 2 & BC-RC > 0.95 | 280 | 169 | 37 | 23 | 14 | 3 | 2 |
| | Undominated | -2 < βNTI < 2 & -0.95 < BC-RC < 0.95 | 2 425 | 1405 | 390 | 285 | 105 | 178 | 63 |

### *Supplementary references*


1. Caporaso JG, Lauber CL, Walters WA, et al (2012) Ultra-high-throughput microbial community analysis on the Illumina HiSeq and MiSeq platforms. ISME J 6:1621–1624. https://doi.org/10.1038/ismej.2012.8

2. Apprill A, Mcnally S, Parsons R, Weber L (2015) Minor revision to V4 region SSU rRNA 806R gene primer greatly increases detection of SAR11 bacterioplankton. Aquat Microb Ecol 75:129–137. https://doi.org/10.3354/ame01753

3. Walters W, Hyde ER, Berg-lyons D, et al (2015) Improved bacterial 16S rRNA gene (V4 and V4-5) and fungal internal transcribed spacer marker gene primers for microbial community surveys. mSystems 1:e0009-15. https://doi.org/10.1128/mSystems.00009-15

4. Hochberg Y (1988) A sharper Bonferroni test for multiple tests of significance. Biometrika 75:800–802. https://doi.org/10.2307/2336325

5. Breiman L (2001) Random Forests. Mach Learn 45:5–32. https://doi.org/10.1023/A:1010933404324

6. Aivelo T, Norberg A, Tschirren B (2019) Bacterial microbiota composition of *Ixodes ricinus* ticks: the role of environmental variation, tick characteristics and microbial interactions. PeerJ 7:e8217. https://doi.org/10.1101/559245

7. Kamvar ZN, Tabima JF, Grünwald NJ (2014) Poppr: an R package for genetic analysis of populations with clonal, partially clonal, and/or sexual reproduction. PeerJ 2:e281. https://doi.org/10.7717/peerj.281

8. R Core Team (2013) R: A language and environment for statistical computing

9. Prevosti A, Ocaña J, Alonso G (1975) Distances between populations of Drosophila subobscura, based on chromosome arrangement frequencies. Theor Appl Genet 45:231–241. https://doi.org/10.1007/BF00831894

10. Cleveland WS, Grosse E, Shyu WM (1992) Local Regression Models. In: Chambers JM, Hastie TJ (eds) Statistical Models in S, 1st ed. Routledge, Boca Raton, LA, USA, p 624

11. Mantel N (1967) The detection of disease clustering and a generalized regression approach. Cancer Res 27:209–220. https://doi.org/10.5694/j.1326-5377.1952.tb83832.x

12. Bogdanowicz D, Giaro K, Wróbel B (2012) TreeCmp: Comparison of trees in polynomial time. Evol Bioinforma 2012:475–487. https://doi.org/10.4137/EBO.S9657

13. Lin Y, Rajan V, Moret BME (2012) A metric for phylogenetic trees based on matching. IEEE/ACM Trans Comput Biol Bioinforma 9:1014–1022. https://doi.org/10.1109/TCBB.2011.157

14. Nei M (1987) Molecular Evolutionary Genetics. Columbia University Press, New York, USA

15. Goudet J. (2005) Hierfstat, a package for R to compute and test hierarchical F-statistics. Mol Ecol Notes 5:184-186. https://doi.org/10.1111/j.1471-8286.2004.00828.x

16. Dray S, Dufour AB (2007) The ade4 package: Implementing the duality diagram for ecologists. J Stat Softw 22:1–20. https://doi.org/10.18637/jss.v022.i04
